# Improving protein tertiary structure prediction by deep learning and distance prediction in CASP14

**DOI:** 10.1101/2021.01.28.428706

**Authors:** Jian Liu, Tianqi Wu, Zhiye Guo, Jie Hou, Jianlin Cheng

## Abstract

Substantial progresses in protein structure prediction have been made by utilizing deep-learning and residue-residue distance prediction since CASP13. Inspired by the advances, we improve our CASP14 MULTICOM protein structure prediction system in the three main aspects: (1) a new deep-learning based protein inter-residue distance predictor (DeepDist) to improve template-free (*ab initio*) tertiary structure prediction, (2) an enhanced template-based tertiary structure prediction method, and (3) distance-based model quality assessment methods empowered by deep learning. In the 2020 CASP14 experiment, MULTICOM predictor was ranked 7^th^ out of 146 predictors in protein tertiary structure prediction and ranked 3^rd^ out of 136 predictors in inter-domain structure prediction. The results of MULTICOM demonstrate that the template-free modeling based on deep learning and residue-residue distance prediction can predict the correct topology for almost all template-based modeling targets and a majority of hard targets (template-free targets or targets whose templates cannot be recognized), which is a significant improvement over the CASP13 MULTICOM predictor. The performance of template-free tertiary structure prediction largely depends on the accuracy of distance predictions that is closely related to the quality of multiple sequence alignments. The structural model quality assessment works reasonably well on targets for which a sufficient number of good models can be predicted, but may perform poorly when only a few good models are predicted for a hard target and the distribution of model quality scores is highly skewed.

## 1 Introduction

Protein structure prediction is to computationally predict the three-dimensional (3D) structure of a protein from its one-dimensional (1D) amino acid sequence, which is much more efficient and cost-effective than the gold-standard experimental structure determination methods such as X-ray crystallography, nuclear magnetic resonance (NMR) spectroscopy, and cryo-electron microscopy (cryo-EM). Computational structure prediction becomes more and more useful for elucidating protein structures as its accuracy improves (Moult et al., 2018). Two kinds of structure prediction methods have been developed: template-based modeling and template-free (*ab initio*) modeling. Template-based modeling (TBM) methods first identify protein homologs with known structures for a target protein and then use them as templates to predict the target’s structure (Källberg et al., 2012; Li & Cheng, 2016). A common approach of identifying homologous templates is based on Hidden Markov Models (Remmert et al., 2012). When no significant known template structures are identified, template-free modeling (FM) is the only viable approach to build structures from protein sequences. Traditional FM methods, such as Rosetta (Rohl et al., 2004), attempt to build tertiary structure by assembling the mini-structures of small sequence fragments into the conformation of the whole protein according to the guidance of statistical energy functions. Other FM tools such as CONFOLD (Adhikari et al., 2015) use inter-residue contact predictions as distance restraints to guide protein folding. In the 13^th^ Critical Assessment of Protein Structure Prediction (CASP13), AlphaFold (Senior et al., 2020), a FM method based on deep learning distance prediction achieved the highest accuracy on both TBM targets and FM targets. Other top CASP13 tertiary structure prediction methods such as Zhang Group (Zheng et al., 2019), MULTICOM (Hou et al., 2019), and RaptorX (Xu & Wang, 2019) were also driven by deep learning and contact/distance predictions.

Inspired by the advances, our CASP14 MULTICOM system is equipped with a new deep-learning based protein inter-residue distance predictor (DeepDist) (Wu et al., 2020) to generate accurate contact/distance predictions, which is used by DFOLD (https://github.com/jianlin-cheng/DFOLD) and trRosetta (Yang et al., 2020) to construct template-free structural models. Moreover, the template-based prediction in MULTICOM is simplified and enhanced by removing redundant template-identification tools and using deeper multiple sequence alignments (MSAs) in template search, while the template libraries and sequence databases are updated continuously. Moreover, 11 new features calculated from predicted inter-residue distance/contact maps are used to predict the quality of protein models in conjunction with other features in DeepRank (Hou et al., 2019) to rank and select protein models.

## 2 Methods

### 2.1 Overview of the MULTICOM system

The pipeline of MULTICOM human and automated server predictors can be roughly divided into six parts: template-based modeling, template-free modeling, domain parsing, model preprocessing, model ranking, and final model generation as depicted in **Figure 1**.

**Figure 1.**
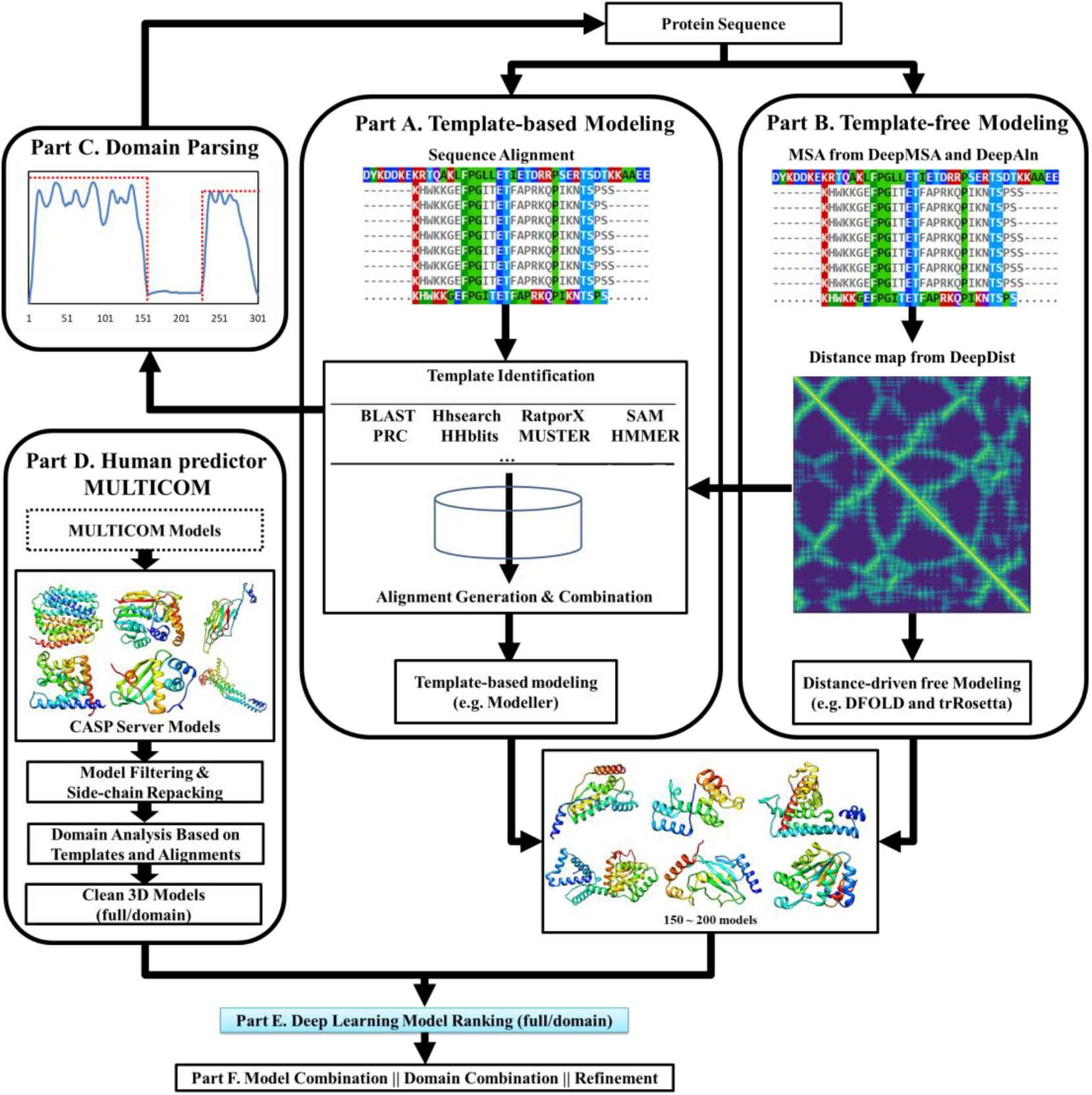
The pipeline of MULTICOM protein structure prediction.

When a target protein sequence is received, template-based modeling (**Part A**) and template-free modeling (**Part B**) start to run in parallel. For the template-based modeling pipeline, MULTICOM first builds the multiple sequence alignments (MSA) for the target by searching it against sequence databases, which are used to generate sequence profiles. Then, the sequences profiles or the target sequence are searched against the template profile/sequence library by various alignment tools (BLAST (Altschul et al., 1990), HHSearch (Söding, 2005), HHblits (Remmert et al., 2012), HMMER (Finn et al., 2011), RatproX (Källberg et al., 2014), I-TASSER/MUSTER (Wu & Zhang, 2008; Yang et al., 2015), SAM (Hughey & Krogh, 1995), PRC (Madera, 2008) and so on to identify templates and generate pairwise target-template alignments. A combined target-template alignment file is generated by combining the pairwise alignments. Structural models are built by feeding the combined alignment file into Modeller (Webb & Sali, 2014). In CASP14, the MULTICOM system was blindly tested as five automated servers. MULTICOM-CLUSTER and MULTICOM-CONSTRUCT servers used the template-based prediction system described above, which was rather slow because it needed to run multiple sequence alignment tools. To speed up prediction, MULTICOM-DEEP and MULTICOM-HYBRID servers only used HHSearch and HHblits in the HHsuite package as well as PSI-BLAST (Altschul et al., 1997) and HMMER to build sequence profiles and search for homologous templates, which are much faster than MULTICOM-CLUSTER and MULTICOM-CONSTRUCT. Considering that the distance-based template-free modeling can often achieve high accuracy on template-based targets, we also tested MULTICOM-DIST server predictor that completely skipped template-based modeling and used only template-free modeling for all the CASP14 targets.

As for the template-free modeling (**Part B**), DeepMSA (Zhang et al., 2020) and DeepAln (Wu et al., 2020) are used to generate two kinds of multiple sequence alignments, which are used to calculate residue-residue coevolution features that are fed into different deep neural networks of DeepDist to predict the distance map - a two-dimensional matrix representing the inter-residue distances for the target protein. For some hard targets, the MSAs generated by HHblits on the Big Fantastic Database (BFD) (Steinegger, Mirdita, & Söding, 2019; Steinegger & Söding, 2018) are also used to predict distance maps. The MSAs along with predicted distance maps are used to generate *ab initio* models with two different *ab initio* modeling tools (e.g., DFOLD and trRosetta (Yang et al., 2020)). In MULTICOM-DEEP and MULTICOM-HYBRID, distance maps and alignments generated by DeepMSA and DeepAln were also used to select templates for template-based modeling.

Domain information (**Part C)** can be extracted from the target-template sequence alignments. If no significant templates are found for a region of the sequence that is longer than 40 residues, the region is treated as template-free (FM) domain, otherwise a template-based domain. And the sequences of the domains are fed into the same pipeline above to build models for individual domains.

For the human predictor (**Part D**), all the CASP server models automatically downloaded from the CASP website and new models generated by MULTICOM servers if any are combined into one model pool as the initial input. Highly similar models from the same groups are filtered out if their pairwise global distance test score (GDT-TS) score is greater than 0.95. SCWRL (Bower et al., 1997) is used to repack the side chains for the models in the filtered model pool. If the target protein is predicted to have multiple domains, the full-length models are split into domain models before model filtering.

Different quality assessment (QA) methods are used in MULTICOM to evaluate the models (**Part E**). For the server predictors in CASP14, the models were assessed by APOLLO (Wang et al., 2011) in MULTICOM-CLUSTER and MULTICOM-HYBRID, by DeepRank (Hou et al., 2019) in MULTICOM-CONSTRUCT, by SBROD (Karasikov et al., 2019) in MULTICOM-DIST, and by the average ranking score of APOLLO, SBROD and distance-based rankings in MULTICOM-DEEP. For the human prediction, three main QA methods are used in model selection, including DeepRank that uses residue-residue contacts predicted by DNCON2 (Adhikari et al., 2018) as input features, DeepRank3_Cluster that uses residue-residue distances predicted by DeepDist as input features, and DeepRank_con that share the same deep network with DeepRank but replace contact predictions from DNCON2 with that from DeepDist. The three QAs also use other features including 1D structural features (e.g., predicted secondary structure, solvent accessibility) and the 3D model quality scores generated by different QA tools (e.g., RWplus (Rykunov & Fiser, 2007), Voronota (Olechnovič & Venclovas, 2014), Dope (Shen & Sali, 2006), and OPUS (Lu et al., 2008)).

Once the QAs generate the model rankings, final models are built by model combination, domain combination or model refinement (**Part F**) from top ranked models. For full-length targets, top five ranked models are combined with other similar top ranked models (maximum 20 models) to generate the consensus models. If a target has multiple domains, top five models are generated by combining domain models using Modeller (Webb & Sali, 2014) or AIDA (Xu et al., 2014). For the human prediction, if the combined models substantially deviate away from the original models, refinement tools (e.g., i3DRefine (Bhattacharya et al., 2016) and ModRefiner (Xu & Zhang, 2011)) will be used instead to refine the top-ranked models to generate the final top five models for submission.

There are several additional differences between the human predictor and server predictors. First, the inputs for the human predictor are the server models from CASP including MULTICOM server models. Additional models generated by MULTICOM servers after the server submission deadline may be added into the model pool for some targets if any. Models filtering and side repacking are applied in the human prediction before feeding the models into the quality assessment methods. Second, in the human predictor, predicted domain boundaries are adjusted based on the top ranked models. Third, in the human prediction, the refinement tools are applied to improve the quality of top ranked models.

### 2.2 Distance-guided template-free modeling

In the distance-guided free modeling, *ab initio* models are mostly generated from predicted residue-residue distance with a customized trRosetta and DFOLD (**Figure 2**). At first, two kinds of MSAs are generated. One is generated by searching a target against the Uniclust30 (Mirdita et al., 2017), UniRef90 (Consortium, 2019) and Metaclust (Steinegger & Söding, 2018) databases using DeepAln and DeepMSA. The other one is generated by using HHblits to search against the BFD (Steinegger et al., 2019; Steinegger & Söding, 2018) database. Then, a combined MSA is built by combining the two kinds of MSAs and filtering out redundancy according to the sequence identity threshold of 95%. The three kinds of MSAs are used for distance prediction and template-free modeling. For two server predictors - MULTICOM-CONSTRUCT and MULTICOM-CLUSTER, the MSAs are then fed into trRosetta to predict the inter-residue geometries and tertiary structures for the target protein. For another three server predictors - MULTICOM-HYBRID, MULTICOM-DEEP, MULTICOM-DIST, the distance maps predicted by DeepDist from the MSAs are used to substitute the default distance maps predicted by the deep networks in trRosetta for model generation. About 50-100 models are built by trRosetta using different probability thresholds on distance map predictions. Top 10 models selected by the ranking methods (e.g., SBROD) are added into the template-free model pool. In these three servers, additional models generated by DFOLD, which takes distance maps predicted from DeepDist as inputs, are also added into the template-free model pool for another round of protein model ranking.

**Figure 2.**
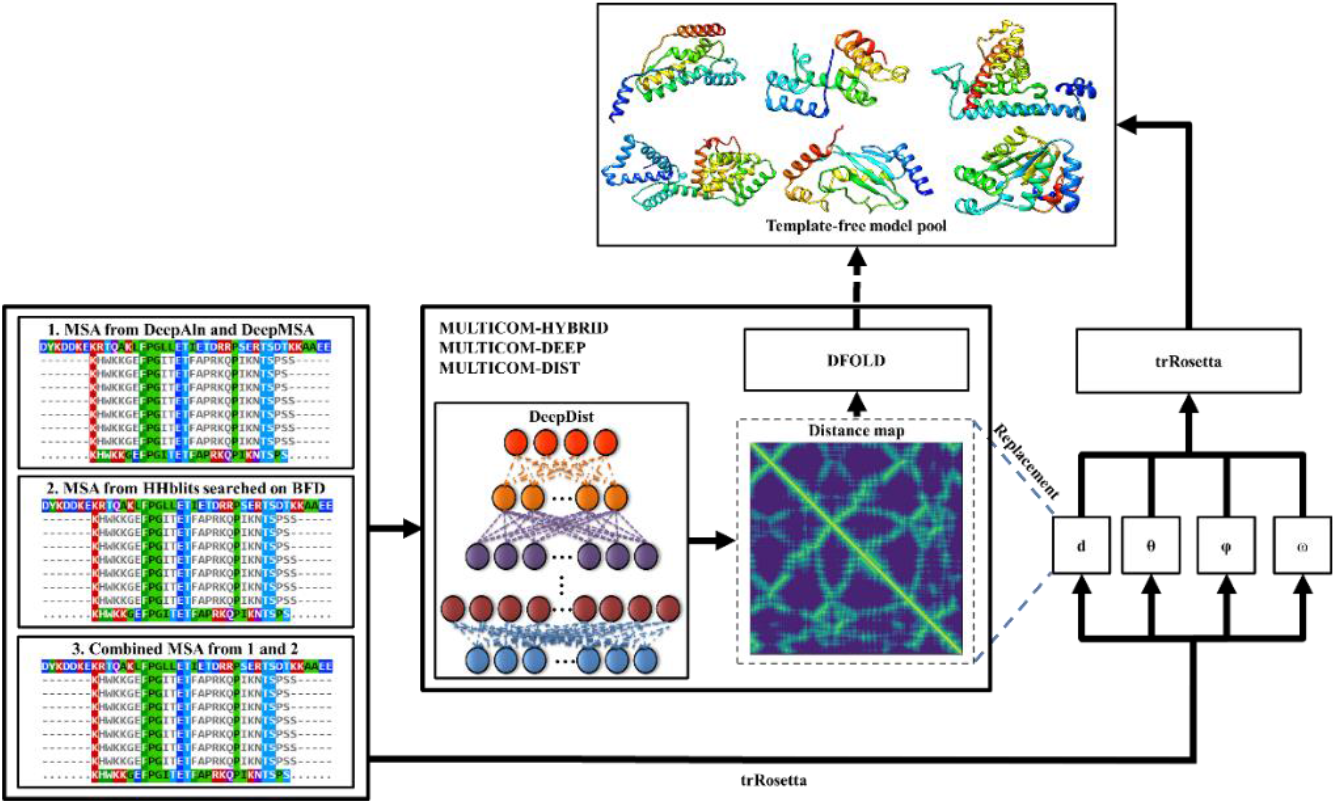
MULTICOM distance-based template-free structure prediction in CASP14.

### 2.3 Protein model ranking

In the MULTICOM human predictor, three main quality assessment (QA) methods (DeepRank, DeepRank_con, DeepRank3_Cluster) are applied to model selection. The methods share the similar features, including 1D features from predicted secondary structures and solvent accessibility and 3D QA scores from different QA tools (i.e., SBROD, RWplus (Rykunov & Fiser, 2007), Voronota (Olechnovič & Venclovas, 2014), Dope (Shen & Sali, 2006), and OPUS (Lu et al., 2008), RF_CB_SRS_OD (Zhang & Zhang, 2010), DeepQA (Cao et al., 2016), ProQ2 (Ray et al., 2012), ProQ3 (Uziela et al., 2016), APOLLO, Pcons (Lundström et al., 2001) and ModFOLDcluster2 (McGuffin & Roche, 2010)), and differ mostly in 2D features derived from predicted contact or distance maps. DeepRank and DeepRank_con share the same neural network and are only different in the input contact map used to generate 2D features. For DeepRank, the input contact map is generated from DNCON2, but DeepRank_con takes an improved contact map from DeepDist as input. For DeepRank3_Cluster, the predicted distance map by DeepDist and the distance map calculated from a 3D model are used to calculate several distance map matching scores (i.e., SSIM & PSNR (Hore & Ziou, 2010), GIST (Friedman, 1979), RMSE, Recall, Precision, PHASH (Niu & Jiao, 2008), Pearson correlation, ORB (Rublee et al., 2011)), which are combined with other 1D and 3D features as inputs. All the quality assessment methods apply the same two-level network architecture. The first level of the network includes 10 neural networks trained by 10-fold cross validation to predict the GDT-TS scores of input models. Then the output scores are combined with initial input features to predict the final scores by the second level network. DeepRank, DeepRank_con and DeepRank3_Cluser were trained and tested on the models of previous CASP experiments before they were blindly applied to the models of the CASP14 experiment.

### 2.4 Model Refinement and Combination

To improve the quality of selected top models, four different methods (e.g., model combination, i3DRefine, ModRefiner, TM-score based combination) are applied under different circumstances in the MULTICOM human predictor. After predicting the quality scores of the input server models, a standard protocol (**Figure 3**) is applied to generate final top 5 models. Each top ranked model is combined with other top ranked models (maximum 20) that are similar to the start model (i.e., GDT-TS > 0.6) to generate a consensus candidate model. If GDT-TS score between the consensus model and the start model is smaller than 0.9, the consensus model is discarded, and the candidate model is generated by using i3Drefine to refine the start model. ModRefiner is used alternatively if severe structural violations (e.g., atom clashes) exist in the candidate model or its secondary structures need to be further improved.

**Figure 3.**
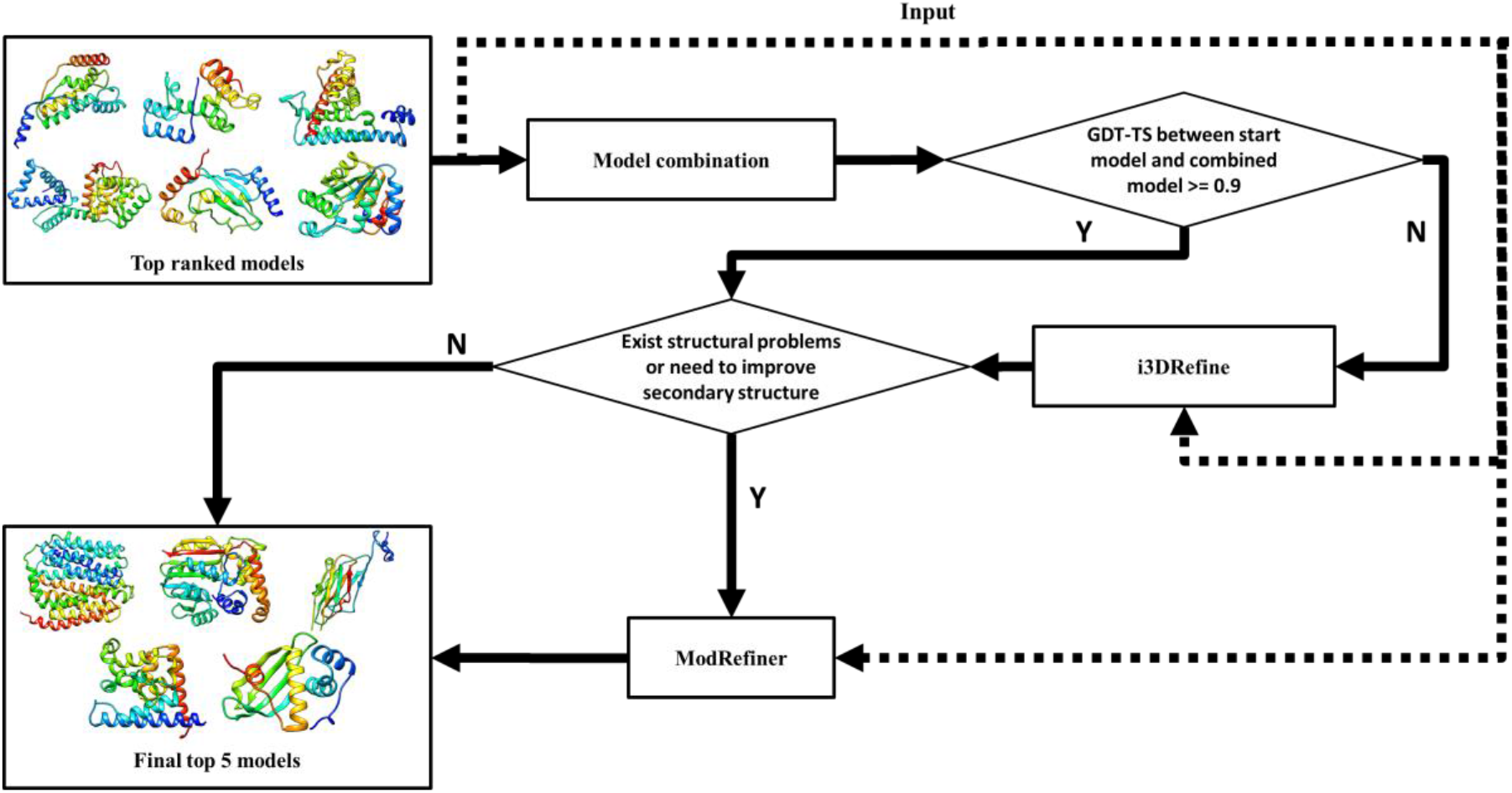
The standard protocol for model refinement and combination.

**Figure 4.**
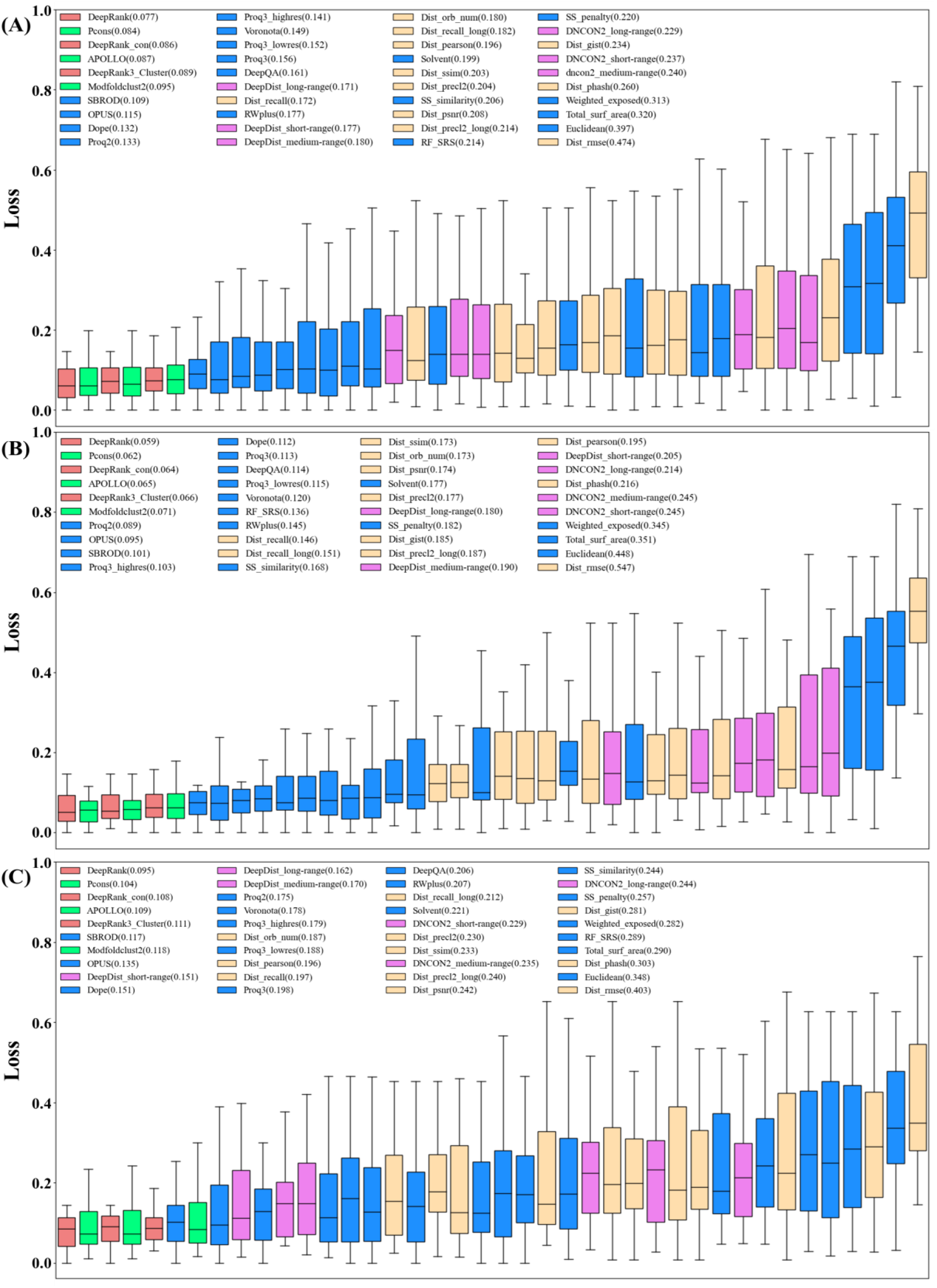
The average loss of 40 QA methods and features in MULTICOM. **(A)** the loss on 61 “all groups” full-length targets. **(B)** the loss on 30 TBM-easy and TBM-hard full-length targets. **(C)** the loss on 31 TBM/FM and FM full-length targets. Red: three DeepRank methods including DeepRank, DeepRank_con, DeepRank3_Cluster; Green: three Multi-model methods including APOLLO (Wang et al., 2011), Pcons (Lundström et al., 2001) and ModFOLDcluster2 (McGuffin & Roche, 2010); Blue: 17 single-model methods including (i.e., SBROD (Karasikov et al., 2019), RWplus (Rykunov & Fiser, 2007), Voronota (Olechnovič & Venclovas, 2014), Dope (Shen & Sali, 2006), OPUS_PSP (Lu et al., 2008), RF_CB_SRS_OD (J. Zhang & Zhang, 2010), DeepQA (Cao et al., 2016), ProQ2 (Ray et al., 2012), ProQ3 (Uziela et al., 2016)); Pink: six contact matching scores including DeepDist/DNCON2 short-range, medium-range and long-range contact matching scores; Yellow: 11 distance scores including SSIM & PSNR (Hore & Ziou, 2010), GIST (Friedman, 1979), RMSE, Recall, Precision, PHASH (Niu & Jiao, 2008), Pearson correlation, and ORB (Rublee et al., 2011).

For some top model, if some of its good regions needed to be kept, but some bad regions needed to be replaced by the corresponding region in another model, a TM-score based model combination method is applied. A superposed model is generated by aligning the two models using TMscore. A preliminary model is generated by replacing the bad region of the top model in the superimposed model with the corresponding region from the other model. The adjusted Ca atom trace of the top model is then extracted from the preliminary model to generate a combined model. The coordinates of other backbone atoms are added into the combined model using Pulchra (Rotkiewicz & Skolnick, 2008). The side chains of the combined model are repacked by SCWRL according to the backbone structure. If needed, ModRefiner is applied to refine the model. This method can also be used to perform domain replacement.

### 2.5 Prediction of the structure for multi-domain proteins

In the MULTICOM system, a domain detection algorithm based on the target-template multiple sequence alignment generated by HHsearch or HHblits is applied to identify domains for multi-domain proteins. Template sequences in the alignment are filtered out by their E-value (> 1), sequence length (<= 40), or alignment coverage (<= 0.5) for the target. If no template is left after filtering, the target is identified as a single-domain template-free target. Otherwise, further analysis is applied to the filtered alignment to identify domains. If a region of the target is not aligned with a template and has more than 40 residues, it is classified as a template-free domain. All the other regions are classified as template-based domains.

After splitting a multi-domain target into domains, the sequence of each domain is fed to the prediction pipeline to generate structural models and top five models are selected. Modeller is used by the default to combine the top domain models into full-length models. AIDA is used alternatively to combine domain models when the full-length model generated by Modeller has severe clashes (i.e., the distance between any two Ca atoms is less than 1.9 Angstrom) or broken chain (i.e., the distance between any two adjacent atoms is greater than 4.5 Angstrom). The domain-based combination models may have good GDT-TS scores for individual domains, but low scores when they are compared with the full-length native structures because they do not have inter-domain interaction information (e.g., relative position and orientation of domains). To address this problem, if a multi-domain target does not have a significant template covering all its domains, domains are treated as independent modeling units and domain-based combination models are used as top prediction. Otherwise, full-length models generated without using domain information are selected based on the domain-based model evaluation to maintain the domain-domain interactions. In some cases, both kinds of models are selected and added into the list of final top five predicted models.

## 3 Results

In CASP14, both MULTICOM human and server predictors participated in the protein tertiary structure prediction. Among 92 CASP14 “all groups’’ domains for tertiary structure prediction, 54 domains are classified as template-based (TBM-easy or TBM-hard) domains that have some structural templates in the Protein Data Bank (PDB) and 38 as FM or FM/TBM domains that have no templates or whose templates cannot be recognized. MULTICOM human predictor was ranked 7^th^ among all the 146 predictors on 92 “all group” domains (https://predictioncenter.org/casp14/zscores_final.cgi). The performance of our human and server prediction methods is systematically analyzed in the following sections using the official evaluation data downloaded from the CASP14’s website.

### 3.1 Performance of MULTICOM human predictor

Based on the official results on the CASP14 website, our MULTICOM human predictor was ranked 7^th^ on all 92 domains overall, 4^th^ on 54 TBM domains and 16^th^ on 38 FM and FM/TBM domains in terms of the sum of the positive Z-scores over the domains. The Z-score of a model predicted for a target is the difference between the GDT-TS score of the model and the average GDT-TS score of all the models predicted for the target divided by the standard deviation of the GDT-TS scores of the models. A positive Z-score indicates that the quality of the model is above the average model. The default CASP14 ranking uses the sum of positive Z-scores over the domains to rank predictors in order not to penalize the new experimental methods that may predict bad models for some targets. Only 7 human predictors from six different groups (AlphaFold2, BAKER, FEIG-R2, Zhang, tFold_human, MULTICOM) achieved higher performance than the best server predictor – QUARK (see **Table 1** for top 20 out 146 predictors and their total Z-score, average TM-score and average GDT-TS score). The average TM-score of MULTICOM on the 92 “all-group” domains is 0.6989, substantially higher than 0.5 – a threshold for a correct fold prediction. If only the top one model per domain is considered, MULTICOM predicts the correct fold for 76 out of 92 (82.6%) domains (i.e., 98% TBM domains and 60.5% FM and FM/TBM domains). If the best of the top five models for each domain is considered, the success rate is increased to 84.8% (i.e., 98% TBM domains and 65.8% FM and FM/TBM domains).

**Table 1.**
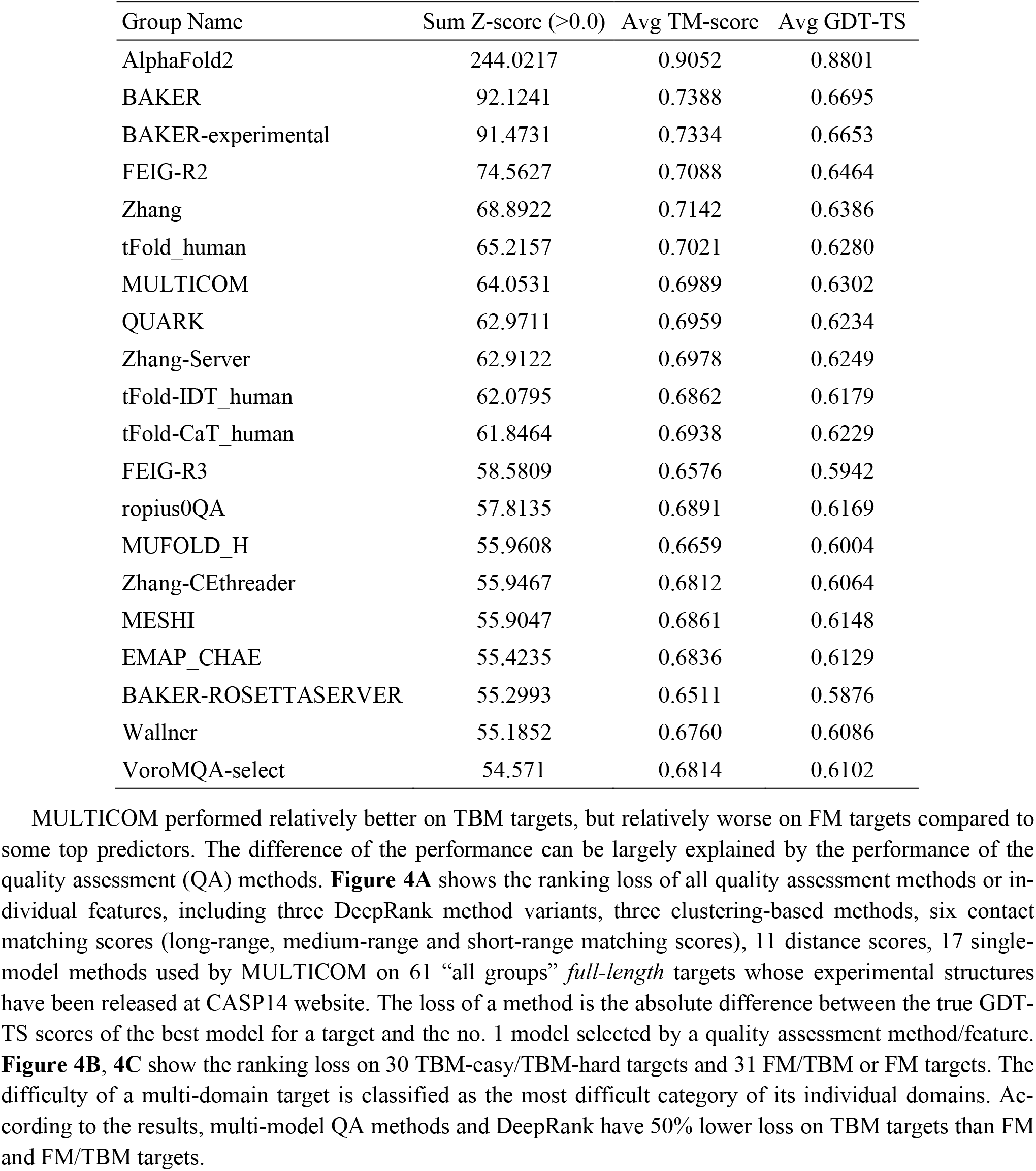
Top 20 predictors in CASP14 tertiary structure prediction.

| Group Name | Sum Z-score (>0.0) | Avg TM-score | Avg GDT-TS |
| --- | --- | --- | --- |
| AlphaFold2 | 244.0217 | 0.9052 | 0.8801 |
| BAKER | 92.1241 | 0.7388 | 0.6695 |
| BAKER-experimental | 91.4731 | 0.7334 | 0.6653 |
| FEIG-R2 | 74.5627 | 0.7088 | 0.6464 |
| Zhang | 68.8922 | 0.7142 | 0.6386 |
| tFold_human | 65.2157 | 0.7021 | 0.6280 |
| MULTICOM | 64.0531 | 0.6989 | 0.6302 |
| QUARK | 62.9711 | 0.6959 | 0.6234 |
| Zhang-Server | 62.9122 | 0.6978 | 0.6249 |
| tFold-IDT_human | 62.0795 | 0.6862 | 0.6179 |
| tFold-CaT_human | 61.8464 | 0.6938 | 0.6229 |
| FEIG-R3 | 58.5809 | 0.6576 | 0.5942 |
| ropius0QA | 57.8135 | 0.6891 | 0.6169 |
| MUFOLD_H | 55.9608 | 0.6659 | 0.6004 |
| Zhang-CEthreader | 55.9467 | 0.6812 | 0.6064 |
| MESHI | 55.9047 | 0.6861 | 0.6148 |
| EMAP_CHAE | 55.4235 | 0.6836 | 0.6129 |
| BAKER-ROSETTASERVER | 55.2993 | 0.6511 | 0.5876 |
| Wallner | 55.1852 | 0.6760 | 0.6086 |
| VoroMQA-select | 54.571 | 0.6814 | 0.6102 |

### 3.2 Performance of MULTICOM in inter-domain structure prediction

In CASP14, MULTICOM was ranked 3^rd^ in a new category (https://predictioncenter.org/casp14/zscores_interdomain.cgi) - inter-domain structure prediction to assess inter-domain interactions for multi-domain targets. The interactions were assessed by Z-score based on F1 score + Z-score based on Jaccard score + Z-score based on best of contact agreement score. 10 multi-domain targets (e.g., T1030, T1038, T1052, T1053, T1058, T1061, T1085, T1086, T1094, T1101) were officially used in evaluation after filtering out targets according to conformational changes, little interaction between domains, and oligomeric interactions. The performance of top 20 out of 136 CASP14 predictors is reported in **Table 2**. MULTICOM’s good performance in this category demonstrates that its modeling strategy for multi-domain targets works well.

**Table 2.** Top 20 predictors in the inter-domain structure prediction.

| Group Name | Sum Z-score (>0.0) | Avg Z-score (>0.0) |
| --- | --- | --- |
| AlphaFold2 | 35.3062 | 3.5306 |
| BAKER-experimental | 15.717 | 1.4288 |
| MULTICOM | 8.986 | 0.8986 |
| BAKER | 8.759 | 0.8759 |
| ProQ3D | 8.5411 | 0.8541 |
| FEIG-R3 | 8.178 | 0.8178 |
| BAKER-ROSETTASERVER | 7.8402 | 0.784 |
| EMAP_CHAE | 7.8057 | 0.7806 |
| VoroCNN-select | 7.5861 | 0.6896 |
| tFold-CaT_human | 7.532 | 0.7532 |
| UOSHAN | 7.2491 | 0.7249 |
| Ornate-select | 7.1811 | 0.6528 |
| Bhattacharya | 7.1549 | 0.7155 |
| ProQ2 | 7.147 | 0.7147 |
| FEIG-R1 | 6.8338 | 0.6834 |
| NOVA | 6.3867 | 0.6387 |
| Bilbul2020 | 6.3768 | 0.6377 |
| RaptorX | 6.3226 | 0.6323 |
| DATE | 6.3098 | 0.631 |

### 3.3 Performance of MULTICOM-CLUSTER, MULTICOM-CONSTRUCT, MULTICOM-HYBRID, MULTICOM-DEEP server predictors using both template-based and template-free modeling

**Figure 5** depicts the performance of the four server predictors on “all group” domains and server-only domains, TBM domains, and FM or FM/TBM domains, respectively. For 92 “all group” and 4 “server only” domains, the average TM-score of the top one models for these targets predicted by MULTICOM-DEEP, MULTICOM-HYBRID, MULTICOM-CONSTRUCT, and MULTICOM-CLUSTER is 0.643, 0.639, 0.640, 0.627, respectively. The average TM-score score of all the servers is substantially higher than 0.5, indicating the MULTICOM servers made good structure prediction for most domains on average. Specifically, if only top one model per domain is considered, MULTICOM-DEEP predicts the correct topology for 75 out of 96 (78.1%) domains (i.e., 55 out of 58 (94.8%) TBM domains and 20 out of 38 (52.6%) FM and FM/TBM domains. **Figure 6** illustrates the predicted structures and distance maps for the 20 FM and FM/TBM domains. If the best of five models for each domain is considered, the success rate is increased to 82.3% for all domains, 98.3% for TBM domains, and 57.9% for FM and FM/TBM domains. Overall, MULTICOM-DEEP performed slightly better than the other three predictors. Their performance of the predictors on the 58 TBM domain is very close, demonstrating that using only HHSearch and HHblits to search for homologous templates in MULTICOM-DEEP and MULTICOM-HYBRID works as well as using multiple alignment and threading tools in MULTICOM-CONSTRUCT and MULTICOM-CLUSTER while substantially reducing the search time. Moreover, MULTICOM-HYBRID and MULTICOM-DEEP performs slightly better on FM or FM/TBM targets than MULTICOM-CONSTRUCT and MULTICOM-CLUSTER, suggesting that replacing the distance maps predicted by trRosetta with the ones predicted by DeepDist may improve the performance of template-free modeling. Furthermore, the average TM-score of all the four predictors on the FM and FM/TBM domains is >= 0.5, substantially better than average 0.32 TM-score of our CASP13 MULTICOM server predictors on the hard domains (Hou et al., 2019), indicating a substantial improvement on template-free modeling made by our new template-free structure prediction method.

**Figure 5.**
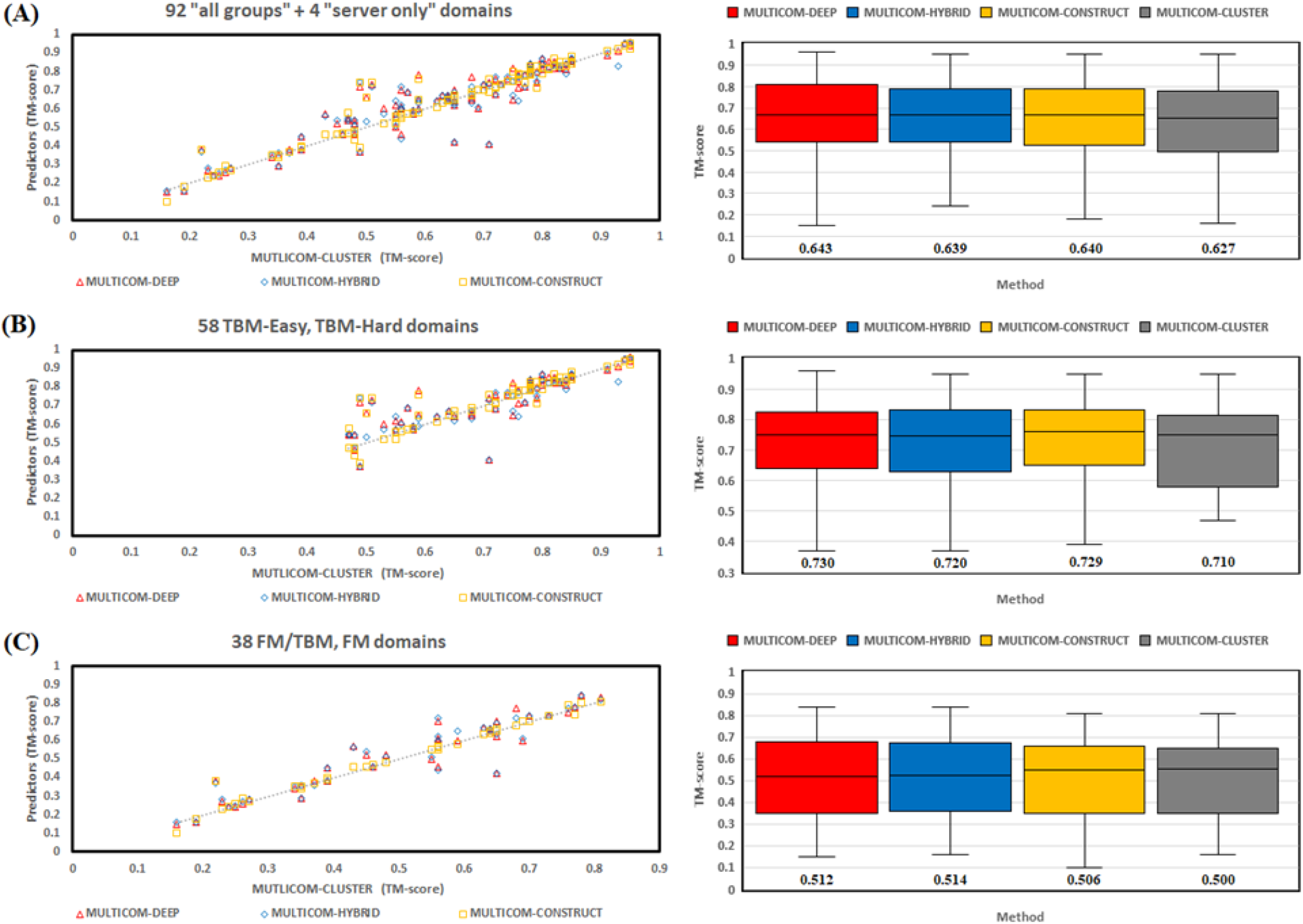
Evaluation of four MULTICOM server predictors in terms of the TM-scores for the first submitted models. **(A)** On 92 “all group” + 4 “server only” domains (left: TM-scores of MULTICOM-DEEP MULTICOM-HYBRID, MULTICOM-CONSTRUCT models vs TM-scores of MULTICOM-CLUSTER models; right plot: mean and variation of the TM-scores of the models of the four methods). **(B)** On 58 template-based (TBM-easy, TBM-hard) domains. **(C)** On 38 FM or TBM/FM domains.

**Figure 6.**
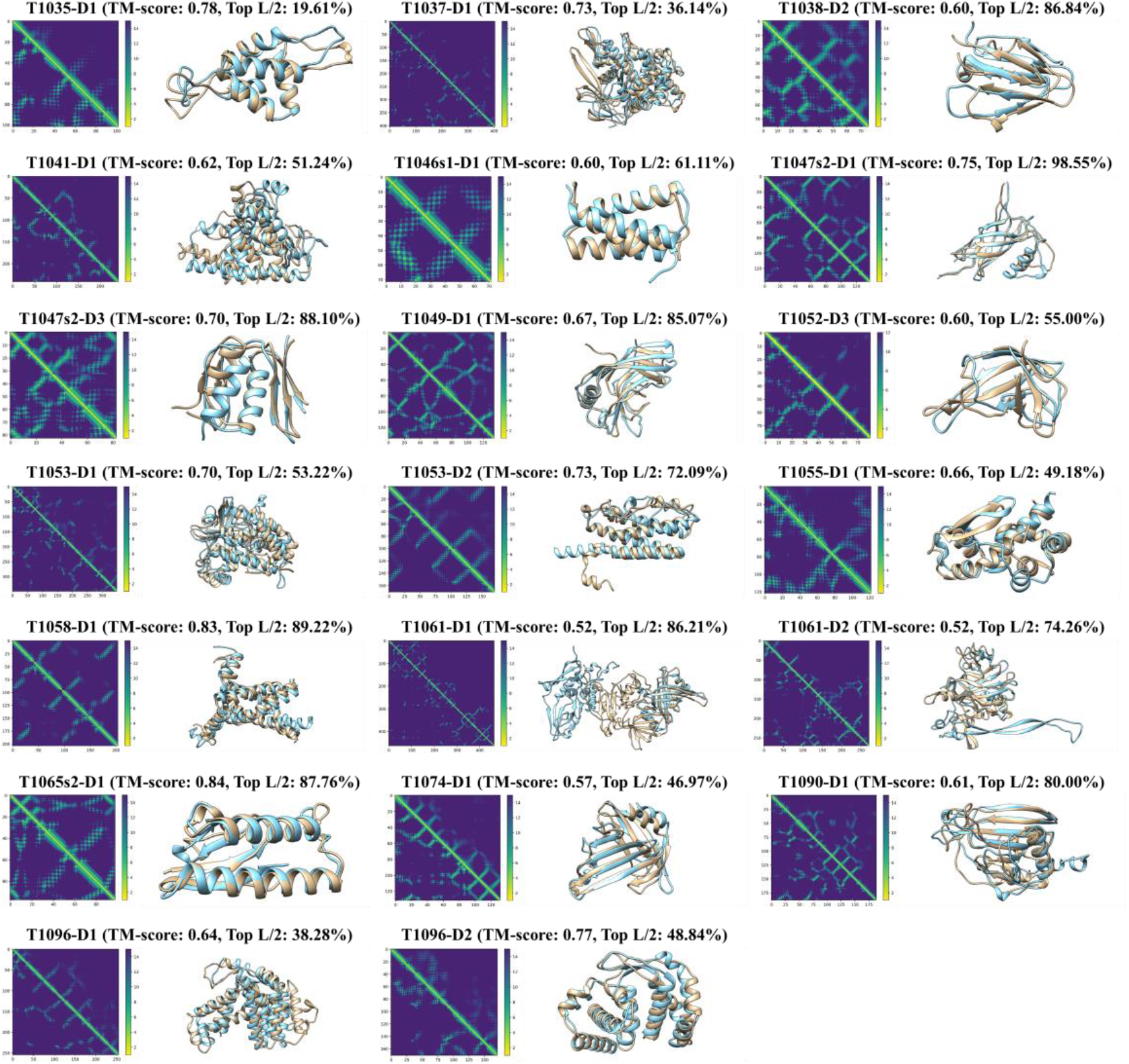
Predicted structures and distance maps compared with native structures and true distance maps for 20 FM or FM/TBM domains for which the first model predicted by MULTICOM-DEEP has the correct topology (TM-score > 0.5). For each domain, on the left is the comparison of the distance maps (lower triangle: true distance map; upper triangle: predicted distance map); and on the right is the comparison of predicted and true structures (light yellow: native structure, light blue: the first predicted structure). The TM-score of the predicted structure and the precision of top L/2 long-range contact predictions for each domain is listed on top of each sub-figure.

### 3.4 Performance of the pure template-free modeling server predictor MULTICOM-DIST

The average TM-score of top one models predicted by MULTICOM-DIST for the 38 CASP14 FM and FM/TBM domains is 0.513, which is similar to 0.514 of MULTICOM-HYBRID or 0.512 of MULTICOM-DEEP. The result is expected because they used the similar distance-based template-free modeling method. Over the 38 CASP14 FM and FM/TBM domains, we investigate how different factors affect the model quality for the distance-based template-free modeling method. One is the number of effective sequences (Neff) in MSAs, measured as the number of the non-redundant sequences at 62% sequence identity threshold. **Figure 7A** shows a weak correlation between the model quality and the logarithm of Neff (Pearson’s correlation coefficient = 0.42) over all 38 domains. But when the logarithm of Neff is less than 6 (i.e., Neff < 400), there is a strong correlation between the model quality and the logarithm of Neff (Pearson’s correlation coefficient is 0.81). The results show the strong positive correlation between the model quality and Neff exists until Neff reaches about 400. Another factor investigated is the precision of distance prediction. **Figure 7B** shows a strong correlation between the precision of top L/2 contact prediction (L: sequence length) and the model quality (Pearson’s correlation coefficient = 0.71), indicating that the model quality increases as the distance prediction gets more accurate.

**Figure 7.**
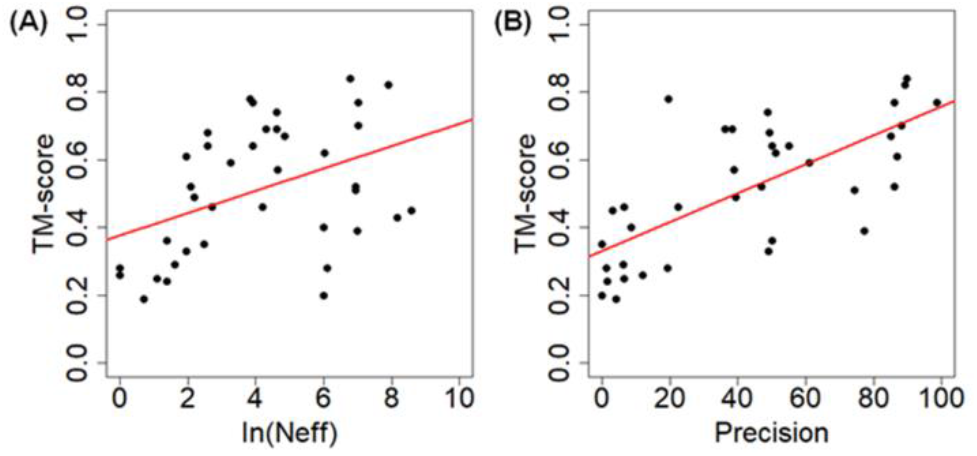
**(A)** Logarithm of Neff of MSA vs. the quality of MULTICOM-DIST top 1 model on the 38 CASP14 FM and FM/TBM domains. **(B)** The precision of top L/2 long-range contact predictions vs. the quality of MULTICOM-DIST top one model on the 38 FM and FM/TBM domains.

On the 58 TBM domains, the average TM-score of MULTICOM-DIST based on template-free modeling only is 0.702, which is a little lower than 0.730 of MULTICOM-DEEP and 0.720 of MULTICOM-HYBRID based on both template-based and template-free modeling. The results show that, even though the template-based modeling may perform slightly better than template-free modeling on some template-based targets, the high TM-score of MULTICOM-DIST on template-based targets and its good performance close to that of MULTICOM-DEEP and MULTICOM-HYBRID demonstrates that the distance-based template-free modeling can work very well on template-based targets, which is consistent with the finding of AlphaFold in the CASP13 experiment. In fact, if only top one model is considered for each domain, MULTICOM-DIST predicts the correct fold for 53 out of 58 (91.4%) TBM domains. If the best of five models is considered for each domain, the success is increased to 96.6%. The results confirm that the distance-based protein structure prediction is able to universally address the protein structure prediction problem. Therefore, the traditional division of protein structure prediction into template-based and template-free modeling may not be necessary anymore, even though template-based structural information can still be used in the modeling process.

The slightly worse average TM-score of MULTICOM-DIST on template-based targets was largely due to the lack of good treatment of large multi-domain targets in the early stage of CASP14 experiment. For large proteins with sequence length > 500, it was often hard to find a sufficient amount of well-aligned homologous sequences covering the entire sequence for accurate full-length residue-residue distance prediction. The global multiple sequence alignment could be dominated by one or two regions with a lot of homologous sequences, leaving the remaining regions not well aligned (i.e., a large number of gaps). For one large TBM-easy target T1036s1 of 818 residues long, MULTICOM-DIST failed to construct the full-length model for this target and its model had a very low TM-score - 0.19 for the domain T1036s1-D1 (sequence region: 1-621). The number of effective sequences of the multiple sequence alignment for the target was 45 and the number of sequences in the multiple sequence alignment was 265, which were relatively small for the distance prediction for the entire target. For each residue position in the multiple sequence alignment of the target, we calculate the number of non-gap amino acids in the position shown in **Figure 8A**. There are few homologous sequences that can cover the entire sequence length. Most homologous sequences in the alignment only cover some regions of the target. There are a large number of gaps in the region ranging from residue 300 to 400. **Figure 8B** compares the true distance map (lower triangle) and the predicted distance map (upper triangle). Even though the predicted distance map contains good intra-domain distance predictions that are similar to the true distances, it does not have good long-range inter-domain distance predictions. The region inside the red circle in the predicted distance map denotes the place where long-range inter-residue contacts were not well predicted in comparison with the true distance map. The true contacts in the region correspond to the interactions between residues 1-78 and residues 57-551 (**Figure 8C**). Different from MULTICOM-DIST, the other four MULTICOM server predictors found strong full-length templates and constructed high-quality models from the templates. For instance, MULTICOM-CONSTRUCT found a significant template 3NWA with the sequence identity of 0.488, sequence coverage of 0.966, and E-value of 5.7E-226 and built a good model with TM-score of 0.92. This example shows that more care needs to be taken for large multi-domain proteins in template-free modeling and it is useful to incorporate some template-based distance information into the distance-based free modeling.

**Figure 8.**
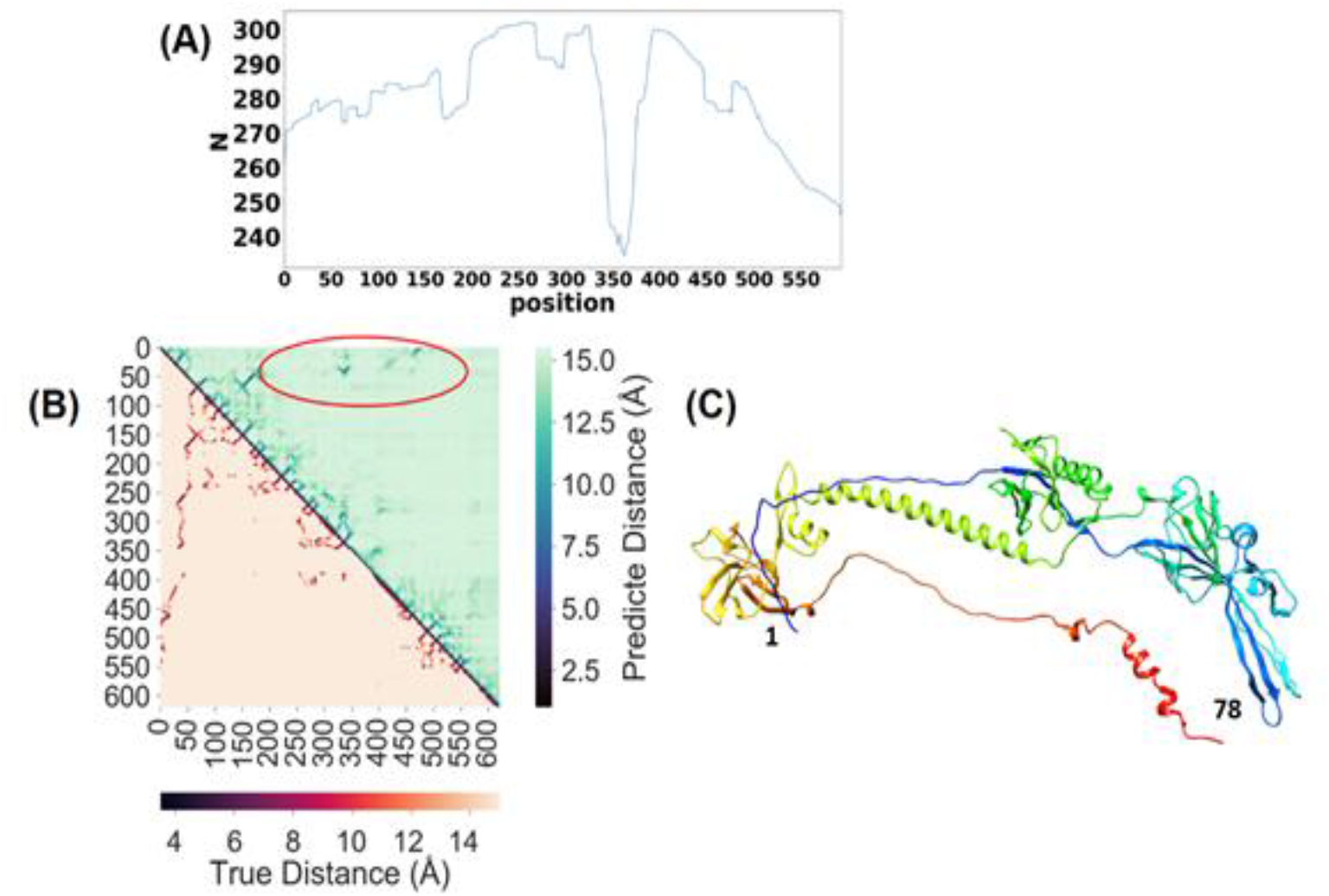
**(A)** The plot of the number of non-gap residues of multiple sequence alignment of T1036s1 against residue positions, where x-axis stands for each residue position and y-axis stands for the number of non-gap amino acids. **(B)** The true distance map of T1036s1-D1 (lower triangle) versus the predicted distance map from MULTICOM-DIST (upper triangle). **(C)** The true structure of target T1036s1-D1 in rainbow, starting from the N-terminal in blue to C-terminal in red.

### 3.5 Comparison of MULTICOM server predictors with other CASP14 server predictors

Based on the official results from the CASP website, after combining multiples server predictors from the same group as one entry, MULTICOM-DEEP was ranked 6^th^ after BAKER, RaptorX, Zhang, FEIG, and Seok groups by the assessor’s formula (GDT_HA + (SG + lDDT + CAD) / 3 + ASE) on 58 TBM domains in terms of sum for Z-scores larger than −2.0 (**Table 3**), where GDT_HA is GDT High Accuracy, SG the Sphere Grinder score, lDDT the local Distance Difference Test score, CAD the Contact Area Difference score, and ASE the Accuracy Self Estimate score. MULTICOM-HYBRID server predictor was ranked 5 ^th^ after Zhang, tFold, BAKER, Yang groups according to the assessor’s formula (GDT_TS + QCS + 0.1 * Molprobity) on 38 TBM/FM and FM domains in terms of the sum for positive Z-scores (**Table 4**), where GDT-TS is the Global Distance Test Score and QCS the Quality Control Score. Both evaluations only considered submitted top 1 models from each server predictor.

**Table 3.**
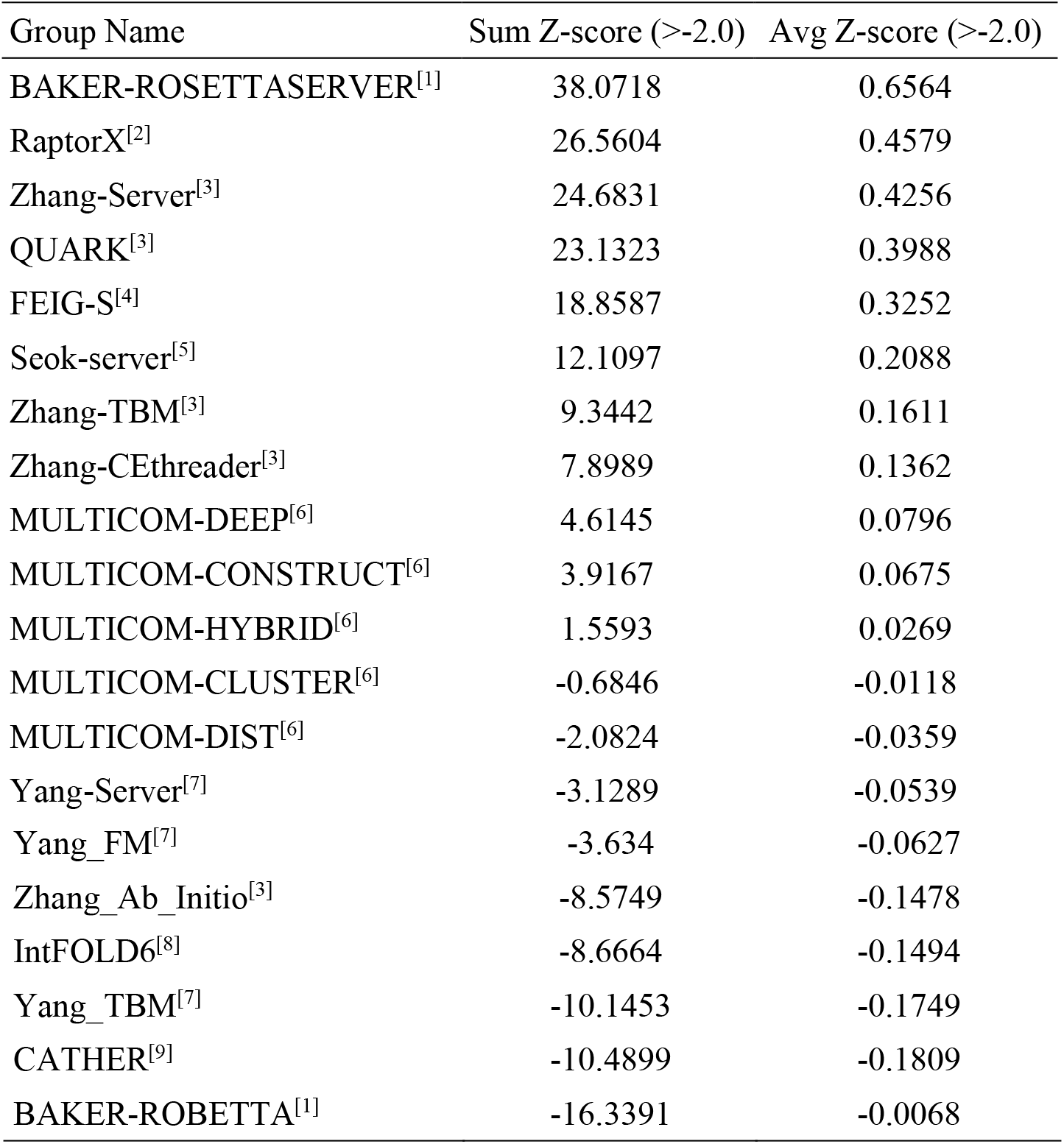
Top 20 CASP14 server predictors on 58 TBM domains. Multiple servers from the same group are denoted by the same number in the superscript.

| Group Name | Sum Z-score (>-2.0) | Avg Z-score (>-2.0) |
| --- | --- | --- |
| BAKER-ROSETTASERVER<sup>[1]</sup> | 38.0718 | 0.6564 |
| RaptorX<sup>[2]</sup> | 26.5604 | 0.4579 |
| Zhang-Server<sup>[3]</sup> | 24.6831 | 0.4256 |
| QUARK<sup>[3]</sup> | 23.1323 | 0.3988 |
| FEIG-S<sup>[4]</sup> | 18.8587 | 0.3252 |
| Seok-server<sup>[5]</sup> | 12.1097 | 0.2088 |
| Zhang-TBM<sup>[3]</sup> | 9.3442 | 0.1611 |
| Zhang-CEthreader<sup>[3]</sup> | 7.8989 | 0.1362 |
| MULTICOM-DEEP<sup>[6]</sup> | 4.6145 | 0.0796 |
| MULTICOM-CONSTRUCT<sup>[6]</sup> | 3.9167 | 0.0675 |
| MULTICOM-HYBRID<sup>[6]</sup> | 1.5593 | 0.0269 |
| MULTICOM-CLUSTER<sup>[6]</sup> | -0.6846 | -0.0118 |
| MULTICOM-DIST<sup>[6]</sup> | -2.0824 | -0.0359 |
| Yang-Server<sup>[7]</sup> | -3.1289 | -0.0539 |
| Yang_FM<sup>[7]</sup> | -3.634 | -0.0627 |
| Zhang_Ab_Initio<sup>[3]</sup> | -8.5749 | -0.1478 |
| IntFOLD6<sup>[8]</sup> | -8.6664 | -0.1494 |
| Yang_TBM<sup>[7]</sup> | -10.1453 | -0.1749 |
| CATHER<sup>[9]</sup> | -10.4899 | -0.1809 |
| BAKER-ROBETTA<sup>[1]</sup> | -16.3391 | -0.0068 |

**Table 4.** Top 20 CASP14 server predictors on 38 TBM/FM and FM domains. Multiple servers from the same group are denoted by the same number in the superscript.

| Group Name | Sum Z-score (>-0.0) | Avg Z-score (>-0.0) |
| --- | --- | --- |
| QUARK<sup>[1]</sup> | 41.0331 | 1.0798 |
| Zhang-Server<sup>[1]</sup> | 40.2236 | 1.0585 |
| Zhang-CEthreader<sup>[1]</sup> | 37.4477 | 0.9855 |
| Zhang-TBM<sup>[1]</sup> | 33.439 | 0.88 |
| Zhang_Ab_Initio<sup>[1]</sup> | 29.5266 | 0.777 |
| tFold-CaT<sup>[2]</sup> | 24.0149 | 0.632 |
| BAKER-ROSETTASERVER<sup>[3]</sup> | 27.0395 | 0.7116 |
| tFold<sup>[2]</sup> | 23.7377 | 0.6247 |
| tFold-IDT<sup>[2]</sup> | 23.2788 | 0.6126 |
| Yang-Server<sup>[4]</sup> | 22.4134 | 0.5898 |
| Yang_FM<sup>[4]</sup> | 21.3196 | 0.561 |
| MULTICOM-HYBRID<sup>[5]</sup> | 20.1767 | 0.531 |
| FoldX<sup>[6]</sup> | 18.905 | 0.4975 |
| MULTICOM-DIST<sup>[5]</sup> | 19.2372 | 0.5062 |
| Yang_TBM<sup>[4]</sup> | 20.0208 | 0.5269 |
| MULTICOM-DEEP<sup>[5]</sup> | 19.6591 | 0.5173 |
| FALCON-DeepFolder<sup>[7]</sup> | 18.6262 | 0.4902 |
| TOWER<sup>[8]</sup> | 18.1948 | 0.4788 |
| MULTICOM-CONSTRUCT<sup>[5]</sup> | 18.4616 | 0.4858 |
| RaptorX<sup>[9]</sup> | 16.2479 | 0.4276 |

### 3.6 Good and bad prediction examples

Among all 92 all-group domains, MULTICOM human prediction were ranked in the top five for three targets in terms of top 1 model: T1034-D1, T1092-D1, and T1093-D2. For T1034-D1 (**Figure 9**), MULTICOM model quality assessment selected RaptorX_TS1 as a start model, whose GDT-TS is 0.8237. MULTICOM combined it with 19 other top ranked server models that were similar to the start model (i.e., GDT-TS > 0.6) to generate a final model. The GDT-TS of the final top1 model (MULTICOM_TS1) is 0.8702, which is significantly improved over the start model and is ranked only after the AlphaFold2 model. For T1092-D1 and T1093-D2, the full-length protein sequences were divided into domains whose boundaries were close to the true domain definition. Based on the domain splitting, MULTICOM was able to select the best domain model in the server model pool as start models to generate high-quality final models.

**Figure 9.**
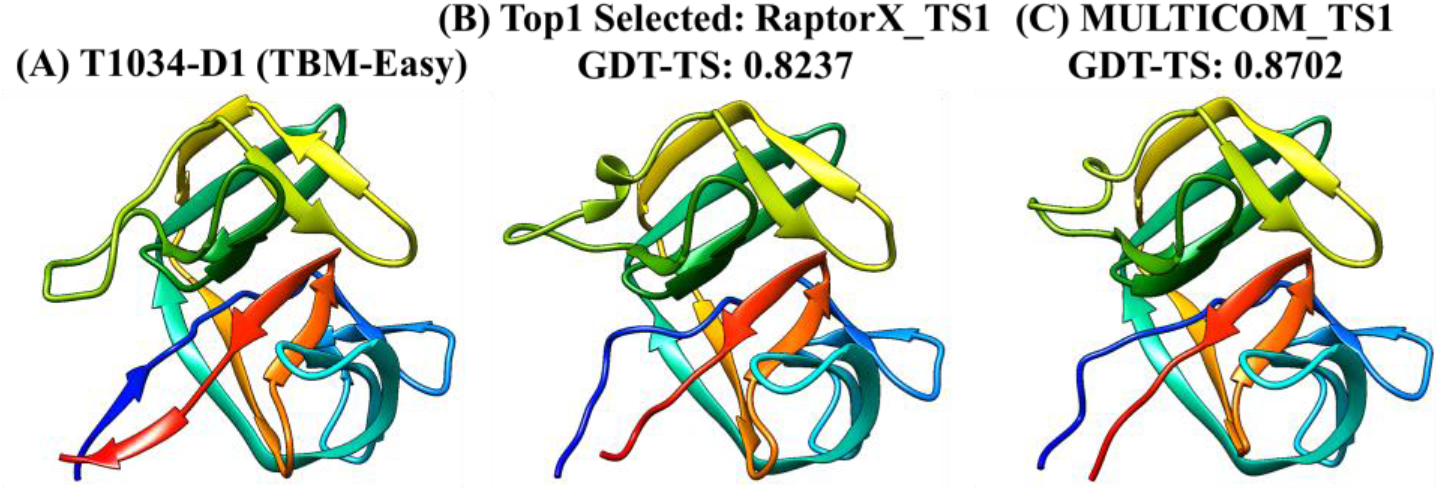
**(A)** The native structure of target T1034-D1; **(B)** the best model selected by the quality assessment; **(C)** the top1 model predicted by MULTICOM.

MULTICOM performed relatively poorly on some FM/TBM or FM domains, including T1031-D1, T1039-D1, T1043-D1, T1061-D1. For T1031-D1, T1039-D1, T1043-D1, MULTICOM’s quality assessment failed to select good start models from the model pool. One reason causing the failure is the number of good-quality models in the model pool is low and the distribution of TM-scores of the models for these targets is highly skewed. In **Figure 10A**, the percentage of good-quality models (TM-score > 0.5) is plotted against the GDT-TS loss of the best quality assessment method - DeepRank. It is shown that TBM targets have a larger proportion of good-quality models than FM and FM/TBM targets. Among five hard targets that have greater than 0% but less than 10% of good models, three of them (T1031-D1, T1039-D1, T1043-D1) have the highest loss among all the targets (>0.25). All the other targets have the loss less than 0.15, even for the targets that have no good models predicted at all (i.e., 0% good models). **Figure 10B** is the plot of the distribution of TM-scores of the models for these three targets. DeepRank selected a model with the score close to the mode (the high density area) of the distribution instead of a good model in the extreme low density area. To further investigate how the distribution of the quality scores of the models in the model pool affects the performance of DeepRank, the skewness of the distribution is calculated for the targets and plotted against the loss on them (**Figure 10C**). The three targets with the highest loss have the highest skewness (i.e., 1.85 for T1031-D1, 1.6 for T1039-D1, 3.05 for T1043-D1), where positive (negative) value of skewness indicates that the mean TM-score is larger (less) than the median TM-score. On 31 FM and FM/TBM targets, the correlation between the skewness and the loss of DeepRank is 0.56, lower than 0.71 of Pcons, indicating that both methods are affected by the skewness, but DeepRank integrating both multi-model and single model features is more robust against the skewness than a clustering-based multi-model method. Another reason for the ranking failure is the incorrect domain prediction. For T1061-D1, a long 949-residue long target, MULTICOM failed to detect the correct domain boundaries, which led to the bad prediction for its first domain (T1061-D1). The example demonstrates that the accuracy of domain prediction has a significant impact on the tertiary structure prediction for some multi-domain targets.

**Figure 10.**
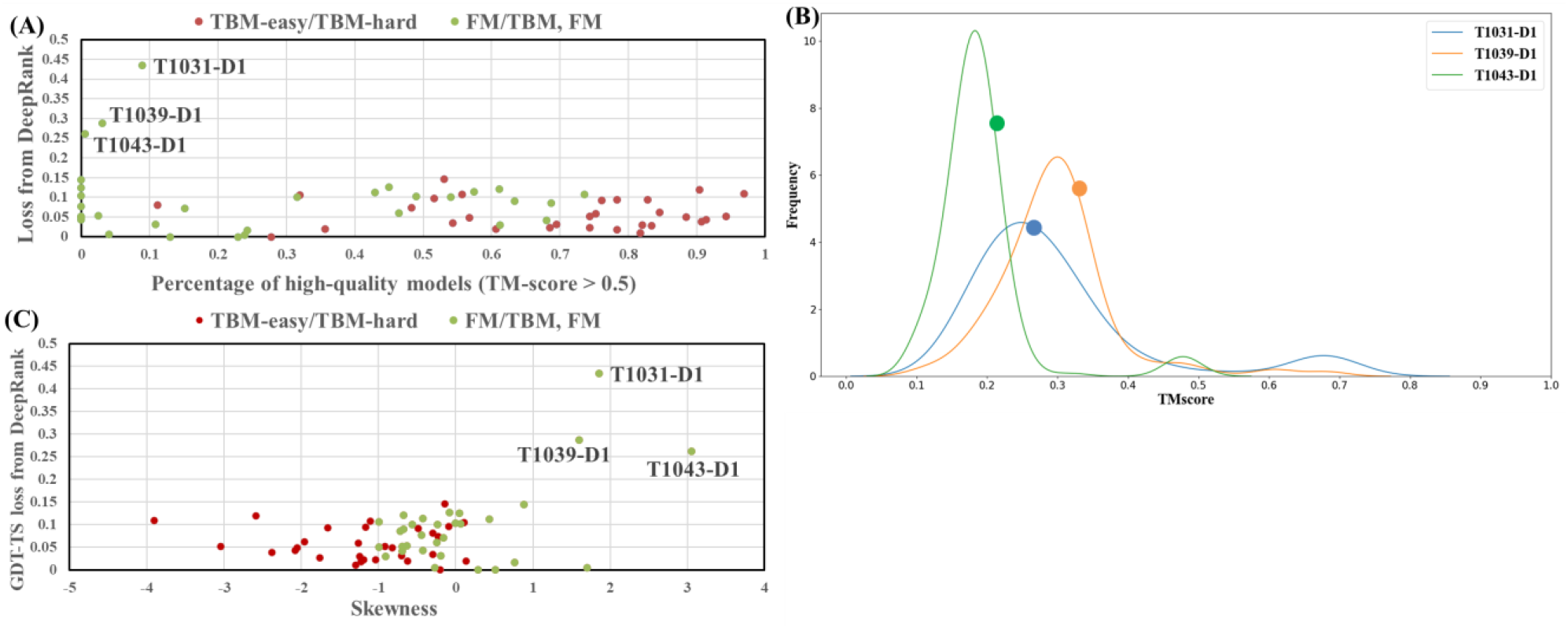
**(A)** The percentage of good-quality models (TM-score > 0.5) vs GDT-TS loss off DeepRank. **(B)** The distribution of TM-scores of the models of T1031-D1 (green), T1039-D1 (red), and T1043-D1 (blue); Dots on the curves denote the top model selected for the targets. **(C)** The skewness of TM-scores of the models vs GDT-TS loss of DeepRank for all 61 targets.

## 4 Conclusion and Future Work

We developed the MULTICOM protein structure prediction system for CASP14 experiment and evaluated and analyzed its performance on CASP14 targets. We demonstrate that the distance-based template-free prediction empowered by deep learning significantly improves the accuracy of protein tertiary structure prediction. The approach can work well on both template-free and template-based targets and therefore can be applied to elucidate the structures of many proteins without known structures in a genome. However, the quality of template-free modeling critically depends on the quality of deep learning-based residue-residue distance prediction, which in turns depends on the quality of multiple sequence alignment. In contrast to the substantial improvement in template-free structure prediction, there is little improvement in protein model quality assessment in our CAS14 system over the CASP13 methods. The quality assessment methods using more accurate residue-residue distance prediction features did not perform better than the quality assessment method using only residue-residue contact prediction features, suggesting that better methods of using distance predictions in quality assessment are needed. Moreover, domain prediction plays an important role in both model generation and evaluation. Accurate domain prediction can help generate better tertiary structure models and select better predicted models for some multi-domain targets. Finally, according to the CASP14 official assessment, our distance-based MULTICOM method that predicts residue-residue distance from multiple sequence alignments first and then reconstructs tertiary structures from the predicted distances did not perform as well as AlphaFold2 that directly predicted 3D structures from multiple sequence alignments, indicating the new direction of the end-to-end prediction of tertiary structures from multiple sequence alignments via deep learning needs to be pursued in the future.

## Acknowledgements

This work is partially supported by Department of Energy grants (DE-SC0020400 and DE-SC0021303), NSF grants (DBI1759934 and IIS1763246), and NIH grants (GM093123).

